# LmbU directly regulates *SLINC_RS02575, SLINC_RS05540* and *SLINC_RS42780*, which are located outside the *lmb* cluster and inhibit lincomycin biosynthesis in *Streptomyces lincolnensis*

**DOI:** 10.1101/2021.06.17.448913

**Authors:** Bingbing Hou, Xianyan Zhang, Yue Mao, Ruida Wang, Jiang Ye, Haizhen Wu, Huizhan Zhang

## Abstract

The productions of antibiotics are usually regulated by cluster-situated regulators (CSRs), which can directly regulate the genes within the corresponding biosynthetic gene cluster (BGC). However, few studies have looked into the regulation of CSRs on the targets outside the BGC. Here, we screened the targets of LmbU in the whole genome of *S. lincolnensis*, and found 14 candidate targets, among of which, 8 targets can bind to LmbU by EMSAs. Reporter assays *in vivo* revealed that LmbU repressed transcription of *SLINC_RS02575* and *SLINC_RS05540*, while activated transcription of *SLINC_RS42780*. In addition, disruptions of *SLINC_RS02575, SLINC_RS05540* and *SLINC_RS42780* promoted the production of lincomycin, and qRT-PCR showed that SLINC_RS02575, SLINC_RS05540 and SLINC_RS42780 inhibited transcription of the *lmb* genes, indicating that all the three regulators can negatively regulate lincomycin biosynthesis. What’s more, the homologues of LmbU and its targets SLINC_RS02575, SLINC_RS05540 and SLINC_RS42780 are widely found in actinomycetes, while the distributions of DNA-binding sites (DBS) of LmbU are diverse, indicating the regulatory mechanisms of LmbU homologues in various strains are different and complicated.

**IMPORTANCE:** Lincomycin is widely used in clinic treatment and animal husbandry. Our previous study firstly demonstrated that LmbU, a novel transcriptional regulator family, functions as a CSR and positively regulates lincomycin biosynthesis. Here, we revealed that LmbU may act as a pleiotropic transcriptional regulator, and directly regulates *SLINC_RS02575, SLINC_RS05540* and *SLINC_RS42780* which are located outside the *lmb* cluster and negatively regulate lincomycin biosynthesis. Interestingly, the homologues of LmbU and its targets are widely found in actinomycetes, indicating the regulatory patterns of LmbU to the targets may exist in a variety of strains. Collectively, our findings elucidated the molecular mechanism with which LmbU regulates the target genes outside the *lmb* culster, and draw a network diagram of LmbU regulation on lincomycin biosynthesis. This lays a solid foundation for the realization of high-yield of lincomycin in industry, and provides the theoretical basis for the functional research of LmbU family proteins.

## Introduction

Streptomycetes are high G+C, filamentous Gram-positive bacteria. In order to cope with the complex and changeable living environment, Streptomycetes evolved a set of protective mechanisms with competitive advantages (1, 2). In this process, a large number of secondary metabolites were produced, including antibiotics with high medical value (3-5). Antibiotic biosynthesis is stringently controlled by precise and pyramidal regulatory cascades (6). *Streptomyces* will monitor the environmental conditions, growing states, population density, and so on, and then secrete and sense specific signal small molecules named autoregulators, including γ-butyrolactones (GBLs), antibiotics and biosynthetic intermediates (7-9). Then, the receptors response and transmit these signal inputs to transcriptional regulators, thereby regulating corresponding antibiotics biosynthesis. Transcriptional regulators in *Streptomyces* are usually classified as global/pleiotropic regulators and CSRs (10). The global/pleiotropic regulators can sense a variety of signals, and not only regulate the biosynthesis of secondary metabolites, but also affect the morphological differentiation of *Streptomyces* (11, 12). The biosynthetic genes of each antibiotic exist in clusters, usually including one or more CSRs, which are at the bottom of the secondary metabolic regulatory network, and directly regulate transcription of the corresponding antibiotics synthetic genes, thus regulating antibiotics biosynthesis (10, 13).

More and more studies have shown that the targets of CSRs are not limited to the gene cluster in which they are situated, but also located in disparate antibiotic biosynthetic gene clusters, forming cross-regulation. For instance, FscRI, a CSR of candicidin gene cluster in *Streptomyces albus* S4, regulates candicidin biosynthesis as well as antimycin biosynthesis (14). In *Streptomyces autolyticus* CGMCC0516, the geldanamycin CSR GdmRIII was found to up-regulate the production of geldanamycin and down-regulate that of elaiophylin by affecting the transcription of the genes in both gene clusters (15). In *Streptomyces venezuelae*, the jadomicin CSR JadR1 can not only activate jadomycin biosynthesis by directly binding to the promoter region of *jadJ*, but also repress chloramphenicol biosynthesis by directly binding to the promoter region of *cmlJ* in chloramphenicol gene cluster (16, 17). Similar examples are also found in coordinated cephamycin C and clavulanic acid biosynthesis by CcaR in *Streptomyces clavuligerus* (18), RED, ACT and CDA biosynthesis by RedZ in *Streptomyces coelicolor* (19), and avermectin and oligomycin biosynthesis by AveR in *Streptomyces avermitilis* (20). Though cases of cross-regulation of disparate antibiotics by one CSR have been reported, screening of the targets of CSRs which located outside the corresponding antibiotics BGSs is barely reported. A case was recently showed that in *Streptomyces cyaneogriseus* ssp. *noncyanogenus*, nemadectin CSR NemR functions as a pleiotropic regulator, which not only activates the transcription of the genes within nemadectin BGC, but also regulates four targets outside the BGC (21). Lincomycin, one of the lincosamide antibiotics, was isolated from a soil-derived Gram-positive bacterium *Streptomyces lincolnensis* in 1962 (22). The 35-kb BGC of lincomycin (*lmb*) contains 25 structural genes (23), three resistance genes (24), and one CSR (25). The structure of lincomycin A is composed of propylproline (PPL) and α-methylthiolincosaminide (MTL), and the biosynthesis of lincomycin comes to light mainly in the recent 10 years (26-31). However, few studies have be reported to elucidate the regulation mechanism of lincomycin biosynthesis. In our previous study, we firstly identified a novel CSR LmbU within lincomycin BGC, and showed that LmbU can positively regulate lincomycin biosynthesis (25). Subsequently, we demonstrated that two global/pleiotropic regulators BldA and AdpA positively regulate lincomycin biosynthesis and morphological differentiation, which function at translational levels and transcriptional levels, respectively (32, 33). Later, Xu et al. showed that a TetR family regulator SLCG_2919 can bind to the promoter regions of lincomycin biosynthetic genes, and directly inhibits lincomycin biosynthesis (34). Li et al. demonstrated that BldD, a famous global regulator, is beneficial to lincomycin biosynthesis and sporulation (35). Xu et al. revealed that a leucine-responsive regulatory protein SLCG_Lrp promotes lincomycin biosynthesis by directly activating transcription of the biosynthetic genes, resistance genes and CSR of lincomycin (36).

In our previous study, we characterized LmbU as a CSR of lincomycin biosynthetic gene cluster, and demonstrated that LmbU homologues are widely found in actinomycetes, indicating LmbU might regulate other target genes except for *lmb* genes. In the present study, we demonstrated that LmbU negatively regulates transcription of *SLINC_RS02575* and *SLINC_RS05540*, while positively regulates transcription of *SLINC_RS42780*. In addition, we showed that *SLINC_RS02575, SLINC_RS05540* and *SLINC_RS42780* can all inhibit the production of lincomycin by repressing transcription of *lmb* genes.

## Materials and methods

### Bacterial strains, plasmids and culture conditions

Bacterial strains and plasmids used in this study are listed in Table 1. *Escherichia coli* strains are and *S. lincolnensis* strains were described in our previous study. Moreover, YEME medium (10 g/L yeast extract, 5 g/L polypeptone, 10 g/L glucose, 3 g/L malt extract, 5 mM MgCl_2_•2H_2_O, 340 g/L sucrose) was used for preparation *S. lincolnensis* mycelium for conjugation, ISP4 medium (10 g/L soluble starch, 1 g/L K_2_HPO_4_, 5 g/L MgSO_4_•7H_2_O, 1 g/L NaCl, 2 g/L (NH_4_)_2_SO_4_, 2 g/L CaCO_3_, 15 g/L Agar, 0.001 g/L FeSO_4_•7H_2_O, 0.001 g/L MnCl_2_•4H_2_O, 0.001 g/L ZnSO_4_•7H_2_O, 0.02 mol/L MgCl_2_) was used for conjugation of *E. coli* and *S. lincolnensis*.

**Table 1.** Strains and plasmids

| Strains or plasmids | Description | Source or reference |
| --- | --- | --- |
| **Strains** |  |  |
| *E. coli* |  |  |
| JM83 | F', ara, $\Delta(lac-pro\ AB)$, *rpsL*, (Str<sup>r</sup>), $\Phi 80$, *lacZ*ΔM15 | (25) |
| ET12567/pUZ8002 | *dam-13::Tn9 dcm-6 hsdM*; containing the non-transmissible RP4 derivative<br>plasmid pUZ8002 | (25) |
| S17-1 | *recA*, *pro*, *hsdR*, RP4-2-Tc::Mu-Km::Tn7 | (37) |
| *S. lincolnensis* |  |  |
| NRRL 2936 | Wild-type (WT), lincomycin producer | NRRL, USA |
| $\Delta lmbU$ | NRRL 2936 with in-frame deletion of *lmbU* | This study |
| WT/p02575TE | NRRL 2936 attBΦC31::p02575TE | This study |
| $\Delta lmbU$/p02575TE | $\Delta lmbU$ attBΦC31::p02575TE | This study |
| WT/p05540TE | NRRL 2936 attBΦC31::p05540TE | This study |
| $\Delta lmbU$/p05540TE | $\Delta lmbU$ attBΦC31::p05540TE | This study |
| WT/p42780TE | NRRL 2936 attBΦC31::p42780TE | This study |
| $\Delta lmbU$/p42780TE | $\Delta lmbU$ attBΦC31::p42780TE | This study |
| $\Delta SLINC\_RS02575$ | NRRL 2936 with in-frame deletion of *SLINC_RS02575* | This study |
| $\Delta SLINC\_RS05540$ | NRRL 2936 with in-frame deletion of *SLINC_RS05540* | This study |
| $\Delta SLINC\_RS42780$ | NRRL 2936 with in-frame deletion of *SLINC_RS42780* | This study |
| **Plasmids** |  |  |
| pLU-1 | *lmbU* cloned in *NdeI/EcoRI* sites of pET-28a (+), for LmbU expression | (25) |
| pKCcas9dO | *actII-orf4*-specific guide-RNA, homologous arms flanking *actII-orf4*,<br>*aac(3)IV*, pSG5 | (54) |
| pKCcas9dlmbU | *lmbU*-specific guide-RNA, homologous arms flanking *lmbU*, *aac(3)IV*,<br>pSG5 | This study |
| pATE152 | pSET152 derivative carrying *xylTE* gene under *lmbAp* promoter | (39) |
| pSET152 | Integrative vector based on $\Phi C31$ integrase | (55) |
| p02575TE | pSET152 carrying *xylTE* gene under the control of *SLINC_RS02575p* | This study |
| p05540TE | pSET152 carrying *xylTE* gene under the control of *SLINC_RS05540p* | This study |
| p42780TE | pSET152 carrying *xylTE* gene under the control of *SLINC_RS42780p* | This study |
| pKCcas9d02575 | *SLINC_RS02575*-specific guide-RNA, homologous arms flanking<br>*SLINC_RS02575*, *aac(3)IV*, pSG5 | This study |
| pKCcas9d05540 | *SLINC_RS05540*-specific guide-RNA, homologous arms flanking<br>*SLINC_RS05540*, *aac(3)IV*, pSG5 | This study |
| pKCcas9d42780 | *SLINC_RS42780*-specific guide-RNA, homologous arms flanking<br>*SLINC_RS42780*, *aac(3)IV*, pSG5 | This study |

### Expression and purification of His_6_-LmbU

The expression plasmid pLU-1 (25) was transformed into *E. coli* BL21 (DE3), and used for His_6_-LmbU expression. The strain was cultivated in 100 mL LB medium at 37 °C until OD_600_ reached about 0.6, 1 mM Isopropyl β-D-1-thiogalactopyranoside (IPTG) was added. After overnight cultivation at 16 °C, the cells were washed twice and suspended in PBS buffer (0.1 M phosphate buffer solution, pH 7.5). Total proteins were released by sonication and His_6_-LmbU was purified using nickeliminodiacetic acid–agarose chromatography (WeiShiBoHui, China). After dialysis and concentration, the purified protein was stored in binding buffer (10 mM Tris-HCl [pH 8.0], 1 mM EDTA, 0.2 mM dithiothreitol, 20 g/ml bovine serum albumin, 1.2% glycerol).

### Electrophoretic Mobility Shift Assay (EMSA)

DNA probes of around 200 bp containing the binding sites of LmbU were amplified via two rounds of PCR. Firstly, primer pairs UBS-X-F/R (X indicates the numbers of the 14 putative targets of LmbU) were used to amplify the cold probes without biotin. Then, biotin-labeled primer EMSA-B* was used for the second-round PCR to generate the labeled probes. The probe prepared by primer pair nag-F/R was used as a negative control. EMSAs were performed as described previously using chemiluminescent EMSA kits (Beyotime Biotechnology, China) with some modification in binding buffer, which included 10 mM Tris-HCl (pH 8.0), 1 mM EDTA, 0.2 mM dithiothreitol, 20 g/ml bovine serum albumin, 1.2% glycerol, and 50 g/ml poly(dI-C) (25). EMSAs performed with 200-fold excesses of specific or nonspecific cold probes were added as controls to confirm the specificity of the band shifts. All primer pairs used in this study are listed in Table S1.

### Construction of *lmbU* disruption strain Δ*lmbU*

To construct a *lmbU* disruption strain, the internal region of *lmbU* (465 bp) was deleted via a CRISPR/Cas9-based genetic editing method (37). The *lmbU*-specific single-molecule-guide RNA (sgRNA) was amplified by PCR using the primer pair sgUF/R with pKCcas9dO as template. Upstream (1.2 kb) and downstream (1.2 kb) homologous arms of *lmbU* were amplified by PCR using primer pairs uU-F/R and dU-F/R, respectively. The *lmbU*-specific deletion cassette was assembled with the above three DNA fragments by using overlapping PCR. Subsequently, the deletion cassette was digested with *Spe*I and *Hin*dIII, and ligated into the corresponding sites of pKCcas9dO. The resulting plasmid pKCcas9dlmbU was introduced into *S. lincolnensis* NRRL 2936 by conjugation, using *E. coli* S17-1 as a donor. The conjugants were selected with nalidixic acid and apramycin, and then identified by PCR using the primer pair JDU-F/R and DNA sequencing. The pKCcas9dlmbU plasmid was eliminated through a few rounds of streak cultivation in YEME medium at 37 °C, which was identified by PCR using the primer pair CR1/2.

### Catechol dioxygenase activity analysis

The regions upstream (relative to the translational start site) of *SLINC_RS02575* (−578 to −1), *SLINC_RS05540* (−471 to −1) and *SLINC_RS42780* (−427 to −1) were amplified using primer pairs p02575-F/R, p05540-F/R and p42780-F/R respectively. The reporter gene *xylTE* was amplified by PCR using primer pair pAxyl-3/4, with pATE152 as a template. Two DNA fragments (promoter region and reporter gene) were cloned into the *Pvu*II site of the integrative plasmid pSET152 using Super Efficiency Fast Seamless Cloning kits (Do Gene, China), resulting in reporter plasmids p02575TE, p05540TE and p42780TE. Then, the reporter plasmids were transferred into the wild-type strain NRRL 2936 and the *lmbU* disruption strain Δ*lmbU*, to construct the reporter strains WT/p02575TE, WT/p05540TE, WT/p42780TE, Δ*lmbU*/p02575TE, Δ*lmbU*/p05540TE and Δ*lmbU*/p42780TE.

Catechol dioxygenase activity analysis was performed as described previously (32). Briefly, the reporter strains were cultivated in YEME medium at 28 °C for one day, then the cells were harvested and lysed by sonication. An appropriate amount of cell extract was added to the assay buffer (100 mM potassium phosphate, pH 7.5, 1 mM catechol), and the optical density at 375 nm was detected per minute. The rate of change per minute per milligram of protein was calculated as catechol dioxygenase activity.

### Bioinformatics analysis (Functional domain analysis, sequence alignment and structure modeling)

Functional domain analysis was performed by BlastP in National Center for Biotechnology Information (NCBI) (https://blast.ncbi.nlm.nih.gov/Blast.cgi?PROGRAM=blastp&PAGE_TYPE=BlastSearch&LINK_LOC=blasthome). Sequence alignment was analyzed using the online software ESPript 3.0 (https://espript.ibcp.fr/ESPript/cgi-bin/ESPript.cgi). Structure modeling was constructed using the online software SWISS MODEL (https://swissmodel.expasy.org/interactive).

### Construction of disruption mutants Δ*02575*, Δ*05540* and Δ*42780*

To construct the *SLINC_RS02575, SLINC_RS05540* and *SLINC_RS42780* disruption strains (Δ*02575*, Δ*05540* and Δ*42780*), the same CRISPR/Cas9-based genetic editing method was carried out as construction of Δ*lmbU* with some modification in construction of disruption plasmids. Take construction of Δ*02575* for example, upstream and downstream homologous arms of *SLINC_RS02575* were amplified by PCR using primer pairs u-02575-F/R and d-02575-F/R, respectively. Specific sgRNA was added to upstream homologous arm by PCR using the primer pair sg-02575/u-02575-R. The above two DNA fragments (sgRNA-containing upstream and downstream homologous arms) were cloned into the *Spe*I/*Hin*dIII sites of pKCcas9dO using Super Efficiency Fast Seamless Cloning kits (Do Gene, China), resulting in disruption plasmid pKCcas9d02575. The primer pair JD-02575-F/R was used to identify the conjugants selected with nalidixic acid and apramycin, and the primer pair CR1/2 was used to verify the elimination of the disruption plasmid.

### Lincomycin bioassay analysis

Lincomycin bioassay analysis was carried out as described in our previous work (25). FM2 medium (20 g/L lactose, 20 g/L glucose, 10 g/L corn steep liquor, 10 g/L polypeptone, 4 g/L CaCO_3_, pH 7.0) was used for fermentation cultivation. *Micrococcus luteus* 28001 was used as an indicator strain, and the concentrations of samples were measured according to the lincomycin standard curves.

Three biological independent experiments were done for the analytical procedures. Error bars indicated means ± standard deviations.

### RNA extraction and quantitative real-time PCR (qRT-PCR)

The strains were cultured in FM2 medium for 2 days, and then RNA was extracted by the method using TRIzol (Thermo Fisher Scientific, United States) (33). The trace amount of DNA was removed through incubation with RNase-free DNase I (TaKaRa, Japan) at 28 °C, and the obtained RNA was analyzed using Nano Drop 2000 (Thermo Fisher Scientific, United States). 1 g RNA was used to synthesize the cDNA using reverse transcription M-MLV (RNase-free) kits (TaKaRa, Japan). qRT-PCR was performed with SYBR green PCR master mix (ToYoBo, Japan) as described previously (25). PCR was carried out in triplicate for each sample. The transcriptional level of *hrdB* was used as a positive internal control to normalize the transcriptional levels of target genes, which were measured by the threshold cycle (2^-ΔΔ*CT*^) method (38).

## Results

### Screening the potential targets of LmbU from the genome of *S. lincolnensis*

Previously, we have identified LmbU as a CSR involved in lincomycin biosynthesis (25), moreover, hundreds of LmbU homologues exist in or outside the BGCs of antibiotics derived from different actinomycetes (39), suggesting that LmbU homologues might not only affect antibiotics biosynthesis by regulating synthetic genes as a CSR, but also participate in other pathways. To explore potential regulatory targets of LmbU in *S. lincolnensis*, a conserved palindrome sequence 5’-TCGCCGGCGA-3’ bound by LmbU was used to scan in the whole-genome of *S. lincolnensis*. A total of 176 conserved sequences were found throughout the genome, among which 54 were located in the potential regulatory regions (–600 – +100 relative to the putative translational start site, and not located inside the operon). Whereafter, 14 candidate targets which may be relevant to lincomycin biosynthesis were selected, including 4 regulators, 5 transporters or resistance related proteins, 2 sigma factors, and 3 other functional proteins (Table 2).

**Table 2.**
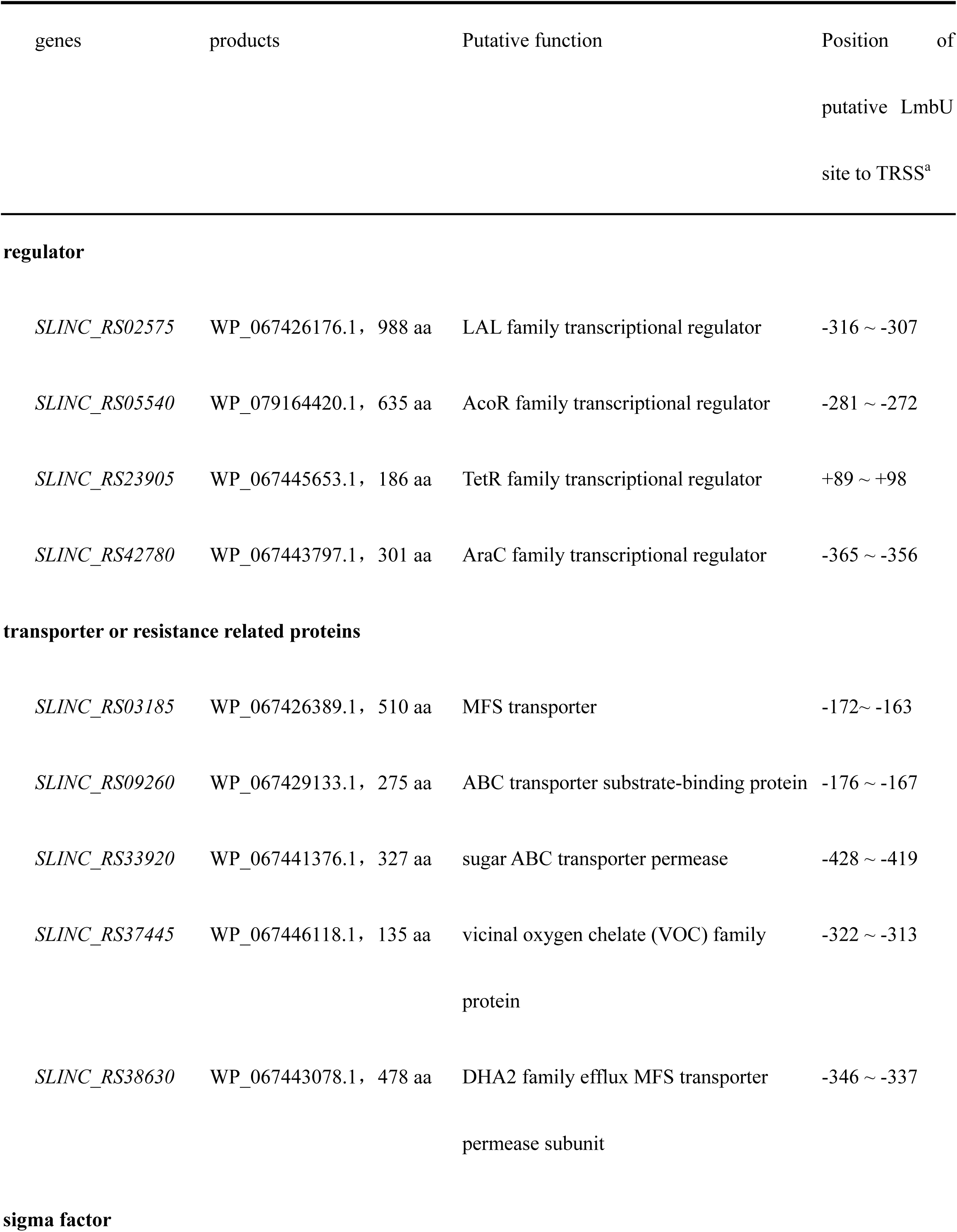
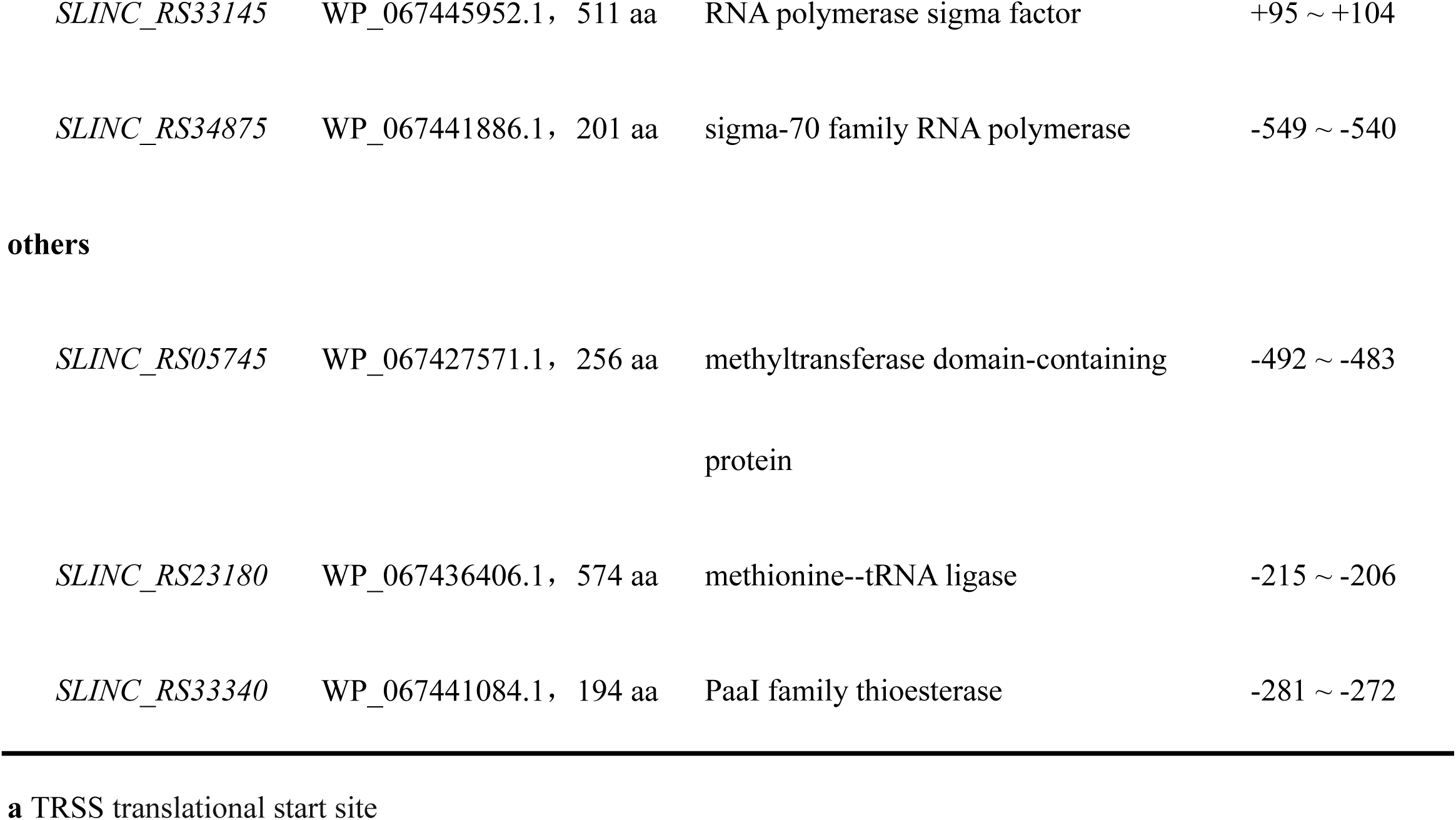
Candidate target genes of LmbU

| genes | products | Putative function | Position of putative LmbU site to TRSS<sup>a</sup> |
| --- | --- | --- | --- |
| **regulator** |  |  |  |
| *SLINC_RS02575* | WP_067426176.1, 988 aa | LAL family transcriptional regulator | -316 ~ -307 |
| *SLINC_RS05540* | WP_079164420.1, 635 aa | AcoR family transcriptional regulator | -281 ~ -272 |
| *SLINC_RS23905* | WP_067445653.1, 186 aa | TetR family transcriptional regulator | +89 ~ +98 |
| *SLINC_RS42780* | WP_067443797.1, 301 aa | AraC family transcriptional regulator | -365 ~ -356 |
| **transporter or resistance related proteins** |  |  |  |
| *SLINC_RS03185* | WP_067426389.1, 510 aa | MFS transporter | -172 ~ -163 |
| *SLINC_RS09260* | WP_067429133.1, 275 aa | ABC transporter substrate-binding protein | -176 ~ -167 |
| *SLINC_RS33920* | WP_067441376.1, 327 aa | sugar ABC transporter permease | -428 ~ -419 |
| *SLINC_RS37445* | WP_067446118.1, 135 aa | vicinal oxygen chelate (VOC) family protein | -322 ~ -313 |
| *SLINC_RS38630* | WP_067443078.1, 478 aa | DHA2 family efflux MFS transporter permease subunit | -346 ~ -337 |
| **sigma factor** |  |  |  |
| *SLINC_RS33145* | WP_067445952.1, 511 aa | RNA polymerase sigma factor | +95 ~ +104 |
| *SLINC_RS34875* | WP_067441886.1, 201 aa | sigma-70 family RNA polymerase | -549 ~ -540 |
| **others** |  |  |  |
| *SLINC_RS05745* | WP_067427571.1, 256 aa | methyltransferase domain-containing<br>protein | -492 ~ -483 |
| *SLINC_RS23180* | WP_067436406.1, 574 aa | methionine--tRNA ligase | -215 ~ -206 |
| *SLINC_RS33340* | WP_067441084.1, 194 aa | PaaI family thioesterase | -281 ~ -272 |

**a** TRSS translational start site

### LmbU binds to the promoter regions of 8 target genes directly

In order to investigate whether LmbU can bind to the above 14 targets, EMSAs were carried out with purified His_6_-LmbU and the DNA probes of candidate targets. The results showed that His_6_-LmbU could obviously bind to the promoter regions of 8 genes in a concentration-dependent manner, but not bind to the promoter regions of other 6 genes (Fig. 1). The deduced products of the 8 target genes were as follows: LAL family transcriptional regulator (encoded by *SLINC_RS02575*), AcoR family transcriptional regulator (encoded by *SLINC_RS05540*), Arac family transcriptional regulator (encoded by *SLINC_RS42780*), MFS transporter (encoded by *SLINC_RS03185*), sugar ABC transporter permease (encoded by *SLINC_RS33920*), DHA2 family efflux MFS transporter permease subunit (encoded by *SLINC_RS38630*), sigma 70 family RNA polymerase (encoded by *SLINC_RS34875*) and methyltransferase domain-containing protein (encoded by *SLINC_RS05745*). Among these, SLINC_RS03185 and SLINC_RS38630 shares 45% identity, and respectively have 36% and 38% identity to LmrA, which is located in the *lmb* cluster and responsible for lincomycin transportation.

**Figure 1.**
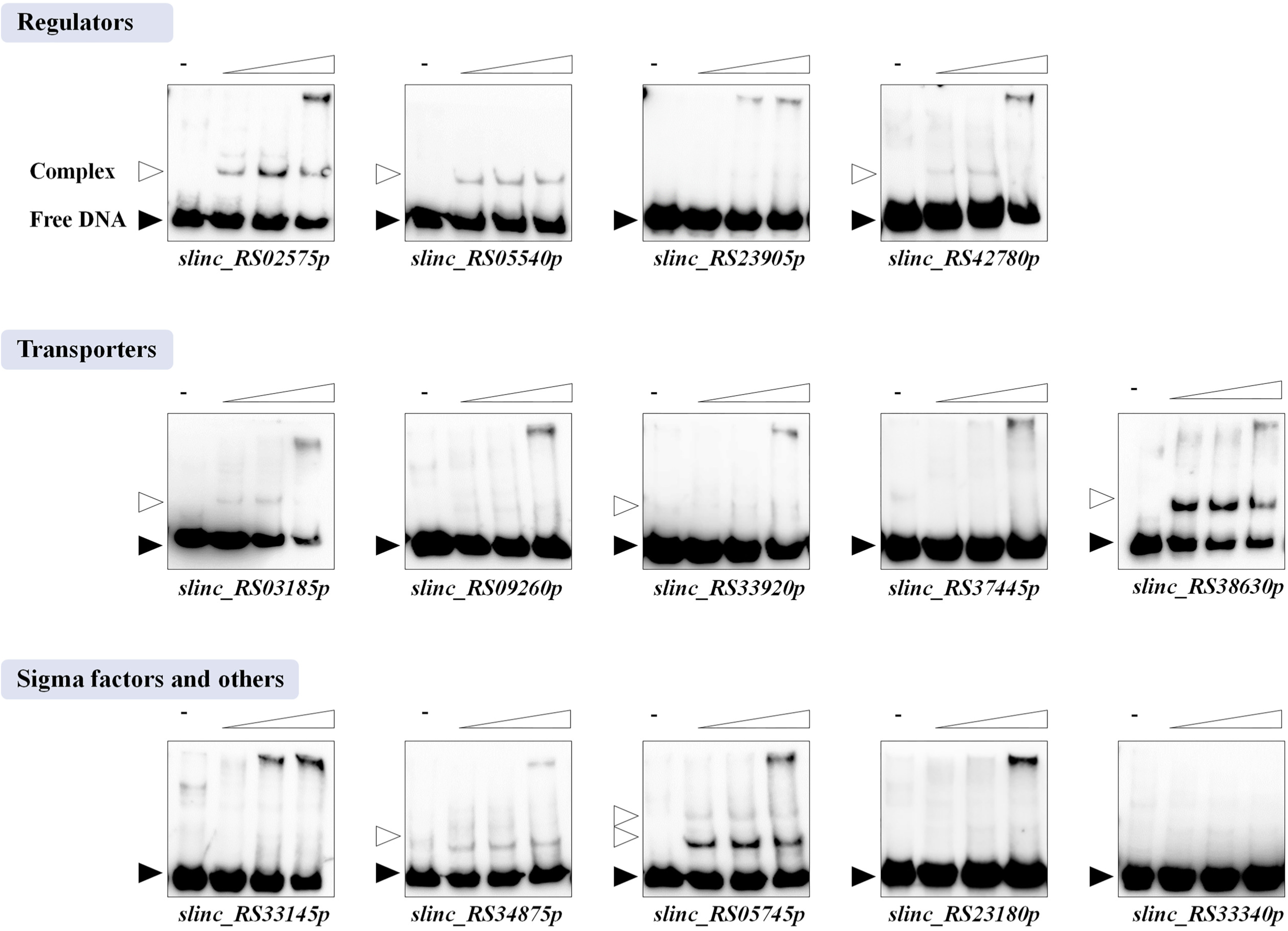
Identification of the binding activities of LmbU to the putative targets. Biotin-labeled probes (5 ng) were incubated with His_6_-LmbU of increasing concentrations (0, 3.2, 6.4 and 9.6 µM). The free probes and DNA-protein complexes are indicated by filled triangles and hollow triangles respectively.

Subsequently, to confirm the binding specificity of His_6_-LmbU to the above 8 targets, competition experiments were introduced into EMSAs. In the presence of 6.4 μM His_6_-LmbU, the retardant bands of all 8 targets were significantly weakened when 200-fold excesses of unlabeled specific DNA were added, but did not change when 200-fold excesses of unlabeled nonspecific DNA (a negative probe that can not bind to His_6_-LmbU, Figure S1) were added. These data demonstrated that LmbU can directly and specifically bind to the promoter regions of the above 8 target genes (Fig. 2), including 3 regulators, 3 transporters, 1 sigma factor and 1 other functional protein. The binding affinities of LmbU with different probes are diverse. LmbU has the highest binding affinity with the probe *SLINC_RS38630p*, while the weakest binding affinity with the probe *SLINC_RS33920p*. Besides, two retardant bands were observed when His_6_-LmbU bound to *SLINC_RS05745p*, indicating the regulatory model of LmbU to this target may be different from that of the others.

**Figure 2.**
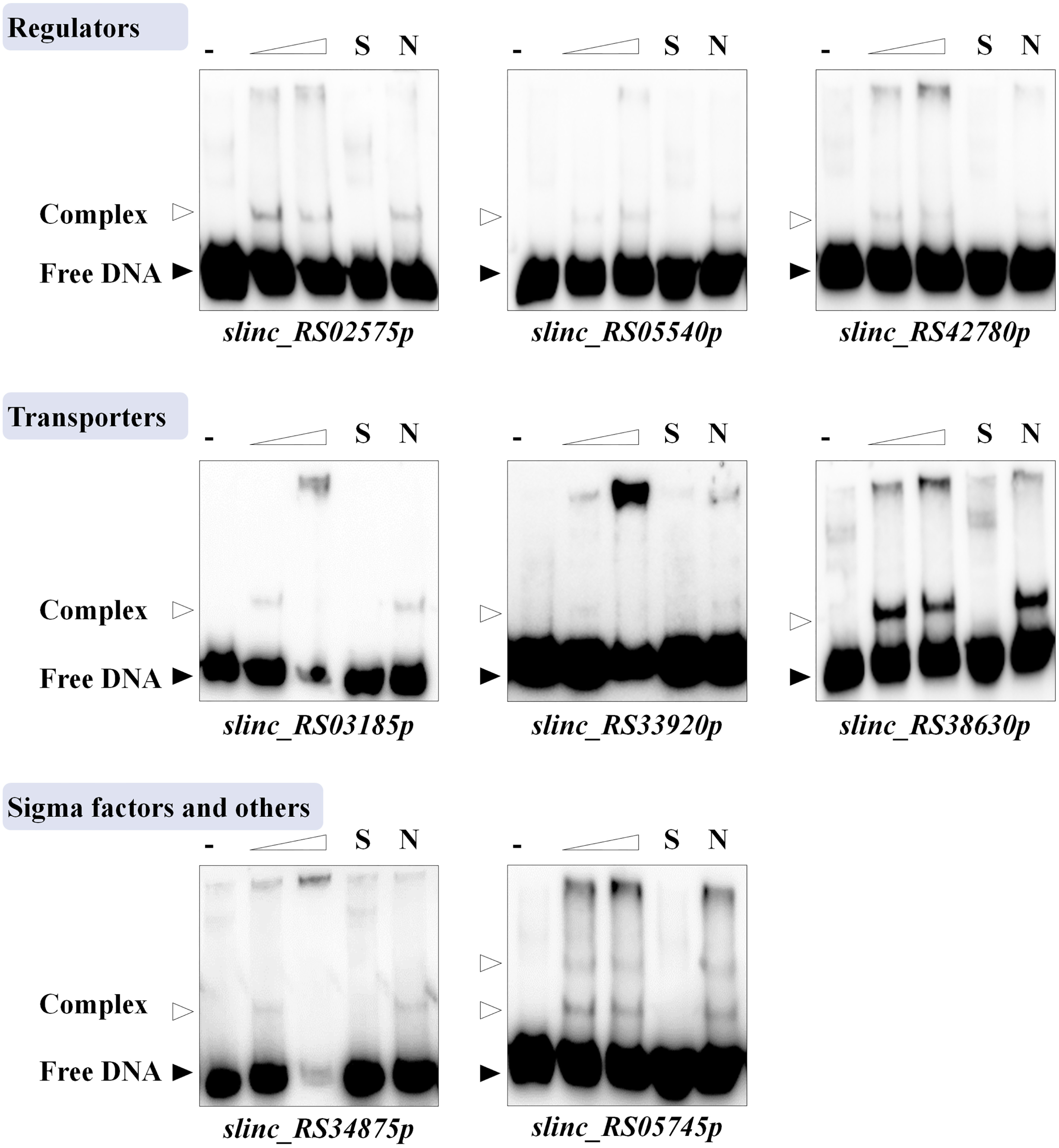
EMSAs of LmbU with promoter regions of target genes. Biotin-labeled probes (5 ng) were incubated with His_6_-LmbU of increasing concentrations (0, 6.4 and 9.6 µM). The free probes and DNA-protein complexes are indicated by filled triangles and hollow triangles respectively. 200-fold excess of specific (S) or nonspecific (N) unlabeled probes were used as competitors of the labeled probes.

As we know, antibiotics biosynthesis is strictly controlled by accurate and sophisticated regulatory networks. Through the above studies, we revealed that three regulatory genes may be regulated by LmbU. Next, our studies will focus on these three regulatory genes.

### LmbU represses the promoters of *SLINC_RS02575* and *SLINC_RS05540* and activates the promoter of *SLINC_RS42780 in vivo*

To investigate the regulation of LmbU to the 3 regulator genes *in vivo*, we firstly constructed a *lmbU* disruption strain Δ*lmbU* by using a CRISPR/Cas9-based genetic editing method. The data of construction and identification of Δ*lmbU* were shown in Figure S2. Then, the WT and Δ*lmbU* strains were chosen for qRT-PCR assays to analyze the effects of LmbU on the transcription of the 3 target genes. However, all the transcriptional levels of the 3 genes were not enough for quantitative analysis (data not shown).

Therefore, we performed *xylTE* reporter assays, using the catechol dioxygenase gene (*xylTE*) as a reporter gene. The reporter plasmids p02575TE, p05540TE and p42780TE, where the *xylTE* gene was under the control of *SLINC_RS02575p, SLINC_RS05540p* and *SLINC_RS42780p* respectively, were introduced into NRRL 2936 and Δ*lmbU*, resulting in reporter strains. As results, enzyme activities of XylTE controlled by *SLINC_RS02575p* and *SLINC_RS05540p* exhibited 7-fold and 6-fold increase in Δ*lmbU* compared to WT, respectively (Fig. 3A and 3B), suggesting that LmbU represses the promoters of *SLINC_RS02575* and *SLINC_RS05540 in vivo*. In contrast, enzyme activities of XylTE controlled by *SLINC_RS42780p* showed 19-fold decrease in Δ*lmbU* compared to WT (Fig. 3C), suggesting that LmbU activates the promoter of *SLINC_RS42780 in vivo*.

**Figure3.**
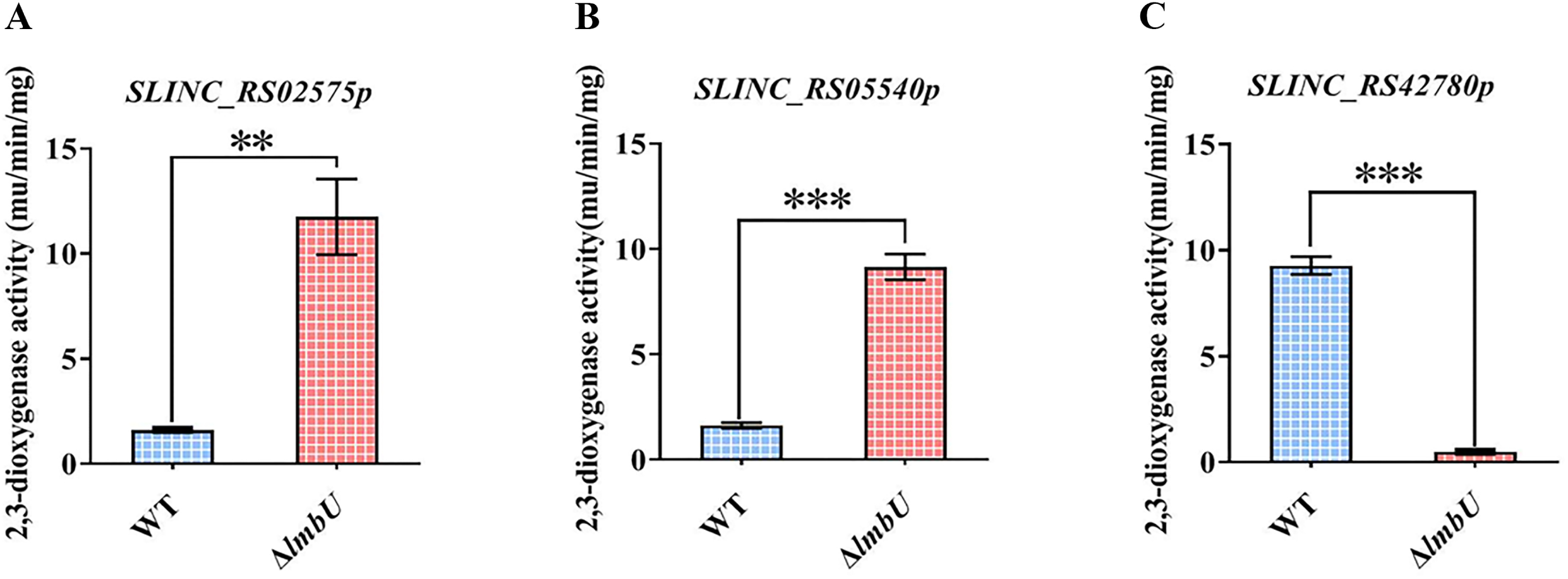
Catechol dioxygenase activity assays of WT and Δ*lmbU* with corresponding reporter plasmids. The results were achieved from three independent experiments. **, *P* < 0.01; ***, *P* < 0.001.

Subsequently, we will further investigate the influence and mechanism of *SLINC_RS02575, SLINC_RS05540* and *SLINC_RS42780* on lincomycin biosynthesis by means of bioinformatics analysis, gene knockout, lincomycin bioassay analysis and qRT-PCR.

### SLINC_RS02575 negative regulates lincomycin biosynthesis

The gene *SLINC_RS02575* is 2967 bp, and encodes a protein containing 988 amino acids, which belongs to a large ATP-binding regulator of the LuxR family (LAL) transcriptional regulator. Sequence alignment showed that N-terminal of SLINC_RS02575 has an AAA+ (ATPases Associated with a wide variety of Activities) domain with ATPase activity, which contains conserved Walker A motif (A/G-X4-G-K-S/T, X indicates any amino acids) and Walker B motif (hhhhhDD, h indicates hydrophobic amino acids) (Fig. 4a and 4b). C-terminal of SLINC_RS02575 has a DNA-binding domain (DBD) of the helix-turn-helix (HTH) structure of the LuxR family (Fig. 4a and 4c). To further investigate the function of SLINC_RS02575 in *S. lincolnensis*, we constructed a *SLINC_RS02575* disruption strain Δ*SLINC_RS02575*, in which the internal region ranging of *SLINC_RS02575* was deleted. The mutant Δ*SLINC_RS02575* was confirmed by PCR using the primer pair JD02575-F/R. PCR products of WT with intact *SLINC_RS02575* gene and Δ*SLINC_RS02575* with defective *SLINC_RS02575* gene were 4.3 kb (Lane 1) and 2.4 kb (Lane 2) respectively. PCR amplification by primer pair CR1/CR2 was used to determine that the disruption plasmid pKCcas9d02575 was eliminated from the mutant Δ*SLINC_RS02575*. A 2.6 kb band appeared only using pKCcas9d02575 as template (Lane 6), rather than WT (Lane 4) or Δ*SLINC_RS02575* (Lane 5). Moreover, sequencing analysis verified that the mutant Δ*SLINC_RS02575* was constructed successfully (Fig. 5a).

Subsequently, the WT and Δ*SLINC_RS02575* strains were cultured in FM2 medium to measure the lincomycin production. *Micrococcus luteus* 28001 was used as an indicator strain to perform lincomycin bioassays. The results showed that the yield of lincomycin in Δ*SLINC_RS02575* increased 2.6-fold compared to that in WT (Fig. 5b), indicating that SLINC_RS02575 negatively regulates lincomycin biosynthesis. Furthermore, qRT-PCR analysis was carried out to assess the influence of SLINC_RS02575 on transcription of *lmb* genes. There are 8 putative operons in the lincomycin cluster, and the first gene of each operon was chosen to perform qRT-PCR, except for *lmbK*, which could not be detected due to the low transcriptional level according to our previous study (25, 33). Compared to WT, the transcriptional levels of *lmbA, lmbD, lmbJ, lmbV, lmbW* and *lmbU* were significantly increased in Δ*SLINC_RS02575* with fold changes 3.7, 2.0, 1.4, 4.3, 13.0 and 3.3, respectively (Fig. 5c). Similar transcriptional levels of *lmbC* were observed in WT and Δ*SLINC_RS02575*, suggesting *lmbC* was not regulated by SLINC_RS02575. These data demonstrated that SLINC_RS02575 can suppress the transcription of *lmbA, lmbD, lmbJ, lmbV, lmbW* and *lmbU*, thereby inhibit lincomycin biosynthesis.

**Figure 4.**
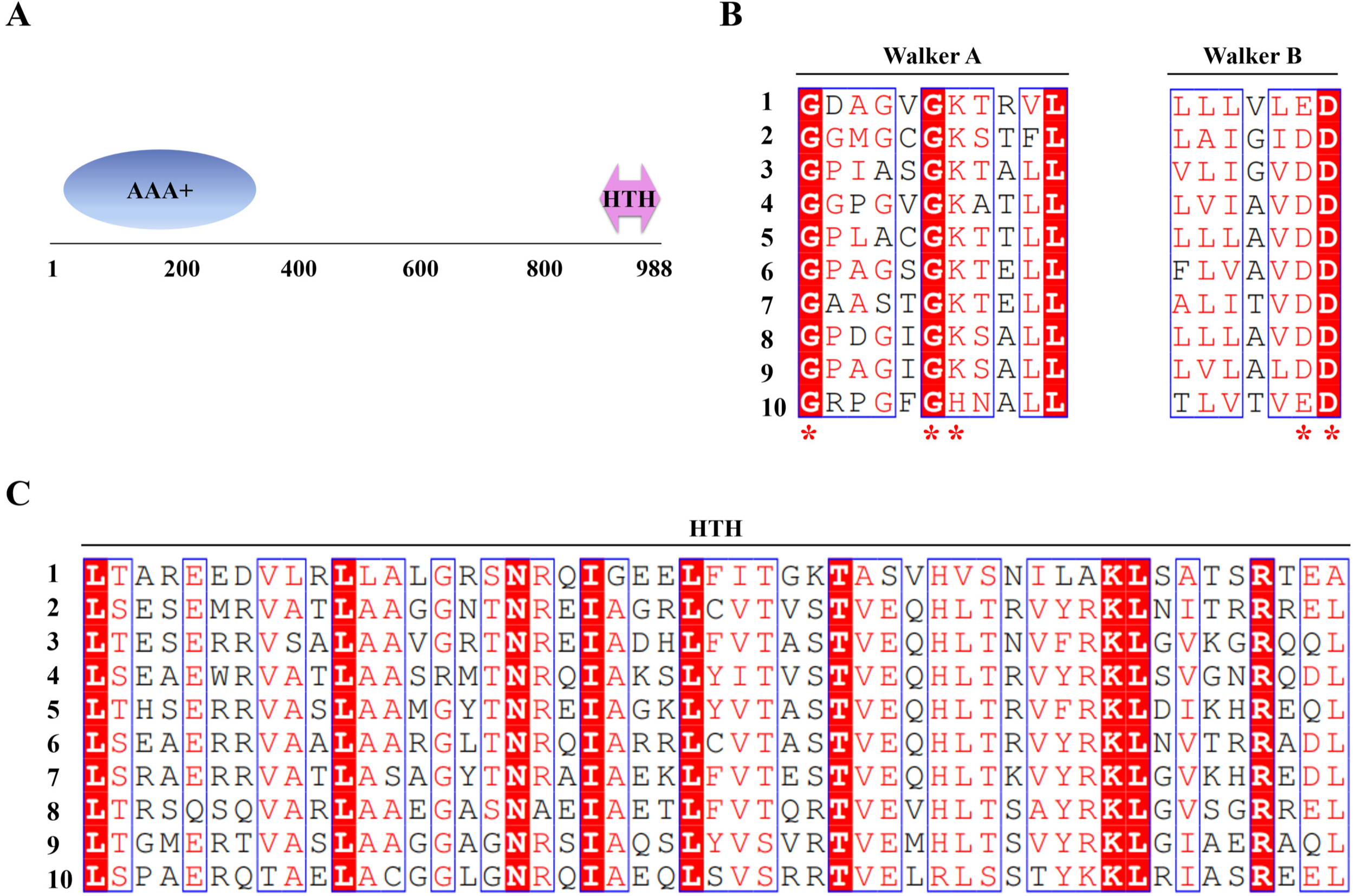
Functional domains and sequence alignment of SLINC_RS02575. **A** Predicted functional domains of SLINC_RS02575. AAA^+^: ATPases Associated with a wide variety of Activities; HTH: helix-turn-helix motif of the LuxR family for DNA binding. **B** Alignment of the AAA domain of SLINC_RS02575 with related proteins. **C** Comparison of the HTH domain of SLINC_RS02575 with that of other proteins. 1, SLINC_RS02575 from *S. lincolnensis* (WP_067426176.1); 2, AveR from *Streptomyces avermitilis* (BAA84600.1); 3, FkbN from *Streptomyces tsukubensis* (TAI41675.1); 4, GdmRI from *Streptomyces hygroscopicus* (ABI93791.1) 5, GdmRII from *Streptomyces hygroscopicus* (ABI93788.1); 6, PikD from *Streptomyces venezuelae* (AAC68887.1); 7, RapH from *Streptomyces hygroscopicus* (AAC38065.1); 8, SalRI from *Streptomyces albus* (ABG02267.1); 9, TtmRI from *Streptomyces ahygroscopicus subsp. wuzhouensis* (AFW98290.1); 10, TtmRII from *Streptomyces ahygroscopicus subsp. wuzhouensis* (AFW98288.1). The conserved amino acids of Walker A and Walker B are indicated by red asterisk.

**Figure 5.**
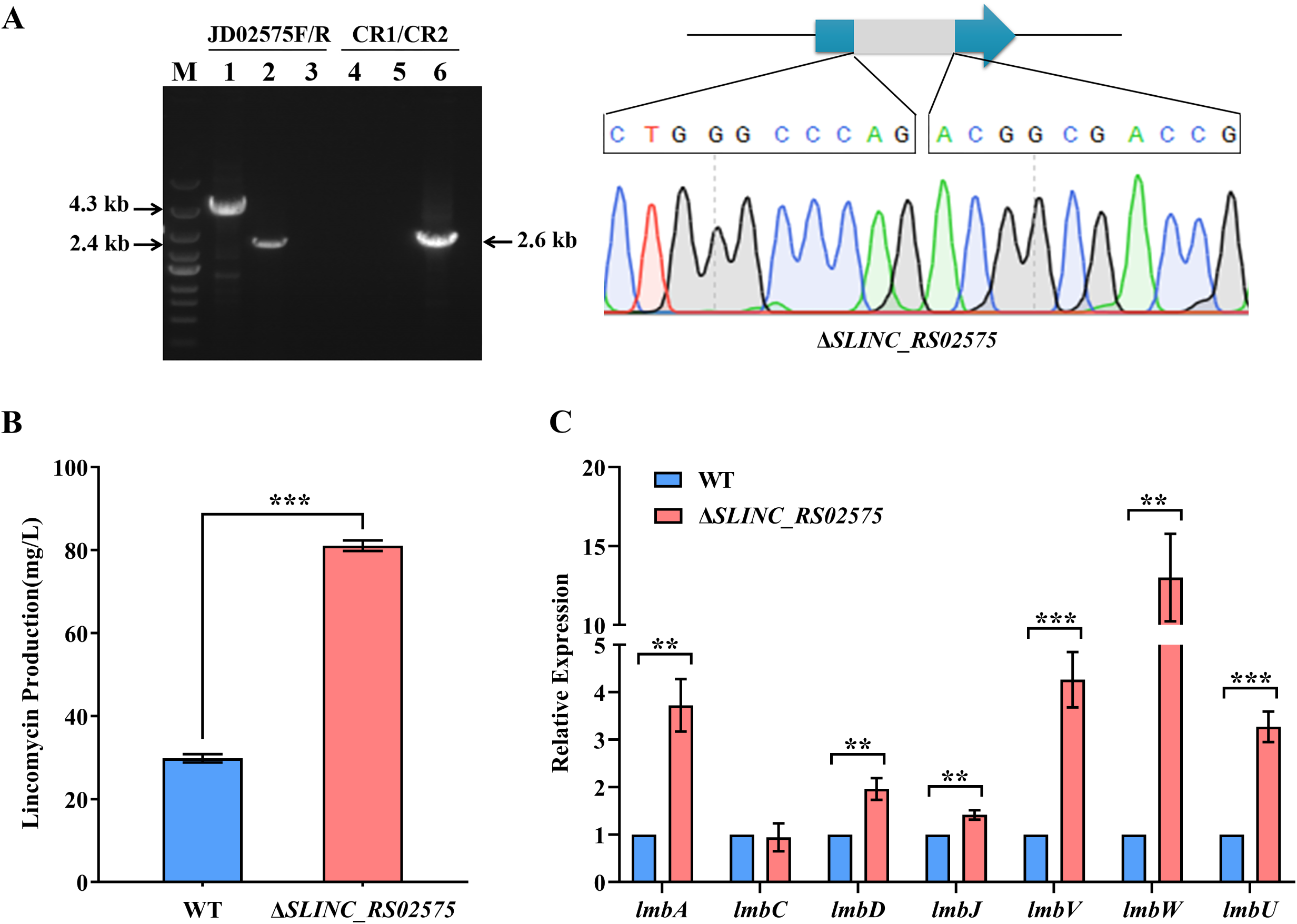
SLINC_RS02575 suppresses lincomycin biosynthesis. **A** Identification of Δ*SLINC_RS02575* by PCR and sequencing. Lane M indicated the DNA molecular weight marker. Lanes 1, 2 and 3 indicated PCR products amplified by primer pair JD02575F/R. Lanes 4, 5 and 6 indicated PCR products amplified by primer pair CR1/CR2. 1 and 4, WT; 2 and 5, Δ*SLINC_RS02575*; 3 and 6, pKCcas9d02575. **B** Effect of SLINC_RS02575 on lincomycin production. **C** Transcriptional analysis of lincomycin biosynthetic genes in WT and Δ*SLINC_RS02575*. The relative expression was normalized using internal reference gene *hrdB*. The transcriptional level of each gene in WT was set to 1.0. **, *P* < 0.01; ***, *P* < 0.001.

### SLINC_RS05540 negative regulates lincomycin biosynthesis

The 1908-bp *SLINC_RS05540* gene encodes an AcoR family transcriptional regulator, which contains 635 amino acids. A conserved Walker A motif (**G**ERGT**GK**) and a HTH motif were found in the internal and the C-terminal of SLINC_RS05540 respectively, indicating SLINC_RS05540 has putative ATP-binding and DNA-binding activities (Fig. 6a). In addition, the results of structure modeling and sequence alignment showed that the HTH motif of SLINC_RS05540 contains three α-helixes referred to the template structure of NtrX derived from *Brucella abortus* (SMTL ID: 5m7n.1, α18-α20) (40), and the amino acids are conserved in *Streptomyces* (Fig. 6b).

**Figure 6.**
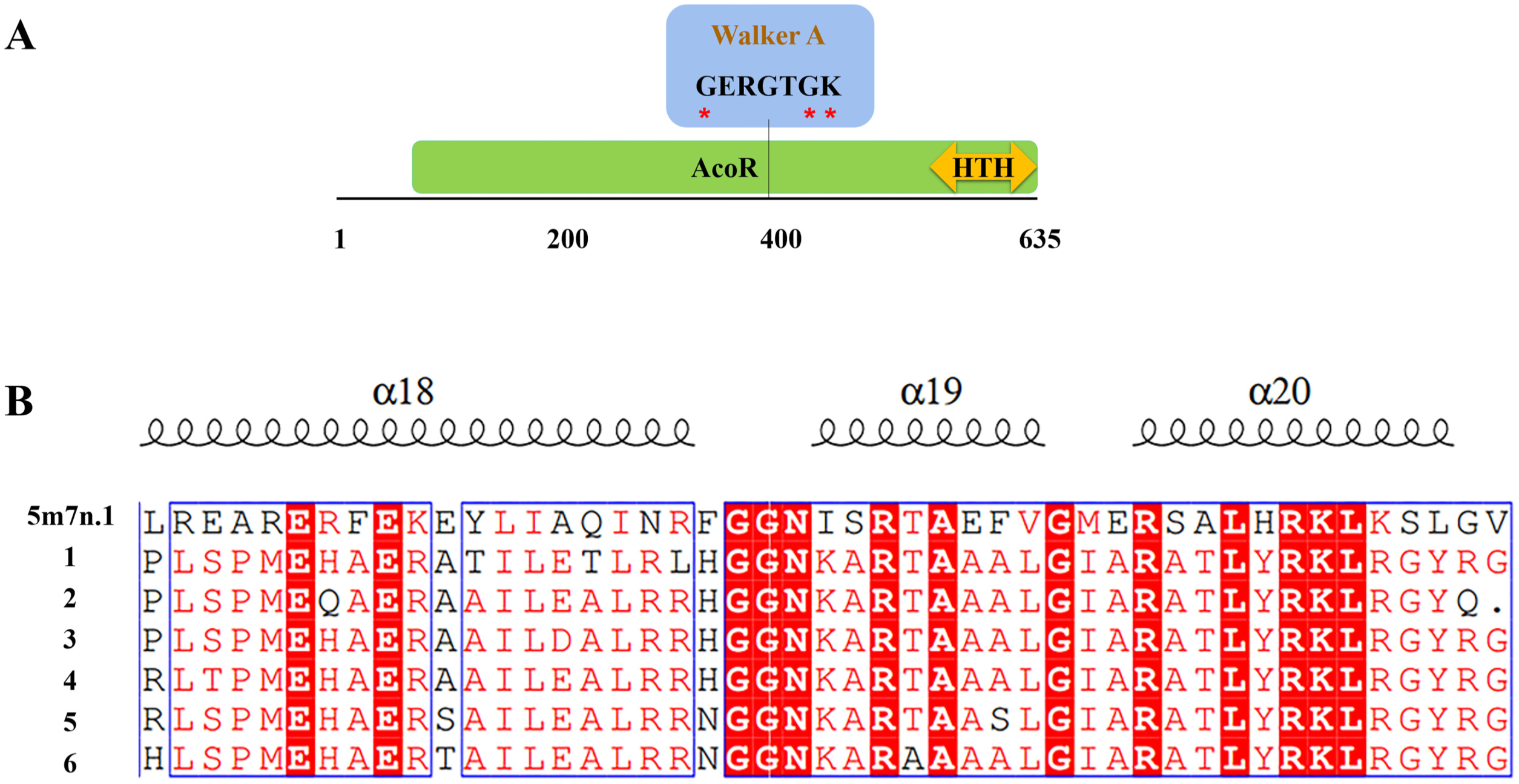
Functional domains and sequence alignment of SLINC_RS05540. **A** Predicted functional domains of SLINC_RS02575. **B** Alignment of the HTH motif of SLINC_RS02575 with that of related proteins. 5m7n.1, the template used for structure modeling of SLINC_RS02575 in SWISS-MODEL. 1, SLINC_RS05540 from *S. lincolnensis* (WP_079164420.1); 2, *Streptomyces aurantiogriseu* (WP_189940635.1); 3, *Streptomyces fulvoviolaceus* (WP_078655870.1); 4, *Streptomyces dysideae* (WP_079085070.1); 5, *Streptomyces scabiei* (WP_037704052.1); 6, *Streptomyces bluensis* (GGZ64430). The conserved amino acids of Walker A are indicated by red asterisk.

Then, a *SLINC_RS05540* disruption strain Δ*SLINC_RS05540* was constructed using the method as above. PCR products amplified by the primer pair JD05540-F/R were 3.8 kb and 2.5 kb with WT and Δ*SLINC_RS05540* as templates, respectively. A 2.6 kb band appeared only using pKCcas9d05540 as template, rather than WT or Δ*SLINC_RS05540* (Fig. 7a). These data indicated that Δ*SLINC_RS05540* was constructed successfully.

**Figure 7.**
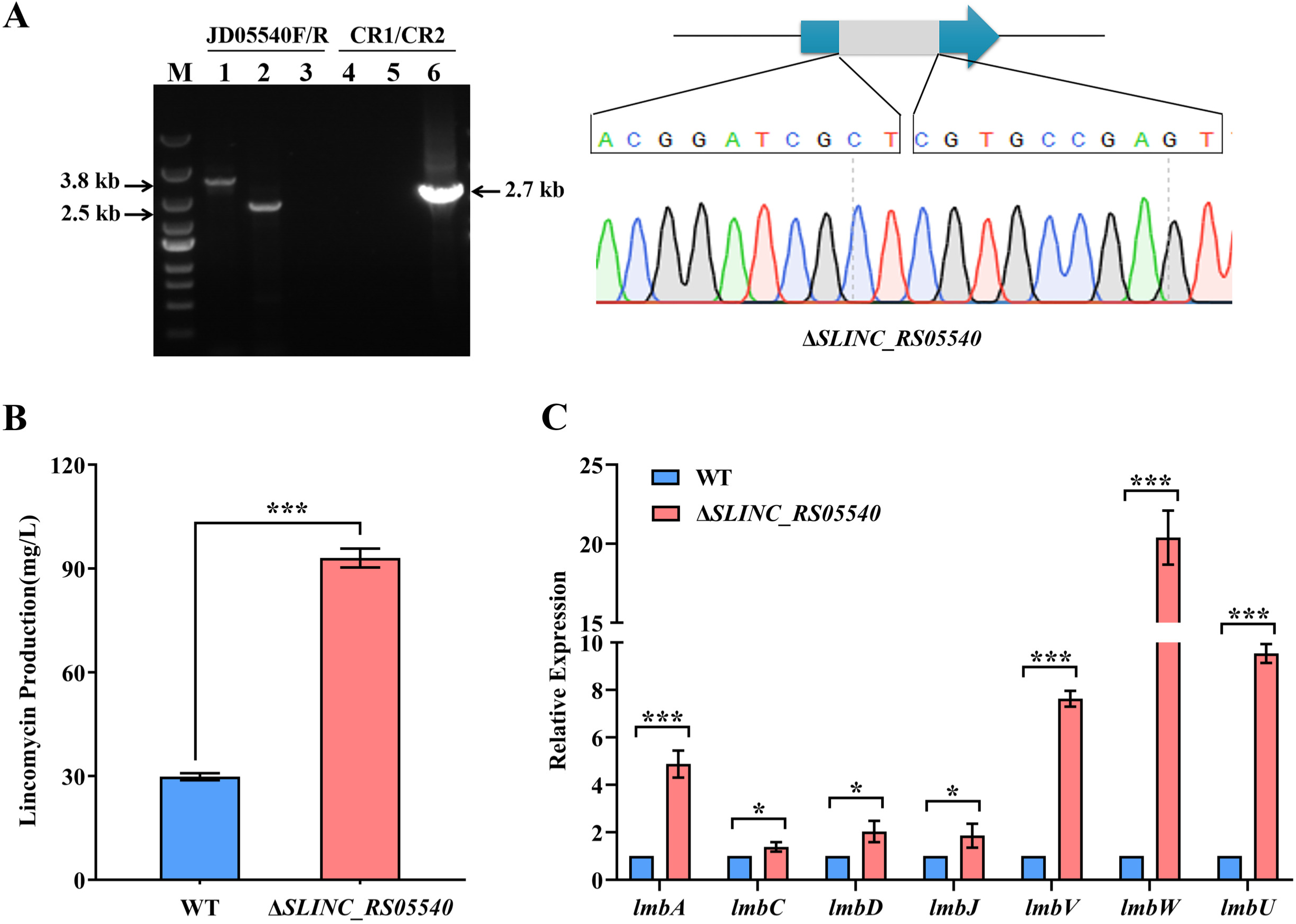
SLINC_RS05540 suppresses lincomycin biosynthesis. **A** Identification of Δ*SLINC_RS05540* by PCR and sequencing. Lane M indicated the DNA molecular weight marker. Lanes 1, 2 and 3 indicated PCR products amplified by primer pair JD05540F/R. Lanes 4, 5 and 6 indicated PCR products amplified by primer pair CR1/CR2. 1 and 4, WT; 2 and 5, Δ*SLINC_RS05540*; 3 and 6, pKCcas9d05540. **B** Effect of SLINC_RS05540 on lincomycin production. **C** Transcriptional analysis of lincomycin biosynthetic genes in WT and Δ*SLINC_RS05540*. The relative expression was normalized using internal reference gene *hrdB*. The transcriptional level of each gene in WT was set to 1.0. *, *P* < 0.05; ***, *P* < 0.001.

Subsequently, the WT and Δ*SLINC_RS05540* strains were cultured in FM2 medium to measure the lincomycin production. The results of lincomycin bioassays showed that the yield of lincomycin in Δ*SLINC_RS05540* increased 3.1-fold compared to that in WT (Fig. 7b), indicating that SLINC_RS05540 negatively regulates lincomycin biosynthesis. qRT-PCR analysis revealed that, compared to WT, the transcriptional levels of *lmbA, lmbC, lmbD, lmbJ, lmbV, lmbW* and *lmbU* were significantly increased in Δ*SLINC_RS05540* with fold changes 4.9, 1.4, 2.0, 1.9, 7.6, 20.4 and 9.5, respectively (Fig. 7c). These data demonstrated that SLINC_RS05540 can suppress transcription of *lmbA, lmbC, lmbD, lmbJ, lmbV, lmbW* and *lmbU*, thereby inhibit lincomycin biosynthesis.

### SLINC_RS42780 negative regulates lincomycin biosynthesis

The 906-bp *SLINC_RS42780* gene encodes a protein containing 301 amino acids, the C-terminal of which contains an AraC type DBD with putative DNA-binding activity (Fig. 8a). SLINC_RS42780 belongs to AraC family transcriptional regulators, which include the famous transcriptional regulator AdpA in *Streptomyces*. It has been reported that the DBD of AdpA derived from *Streptomyces griseus* contains two HTH motifs, HTH1 (α2 and α3) and HTH2 (α5 and α6) (41). Structure modeling and sequence alignment revealed that the DBD of SLINC_RS42780 includes similar HTH motifs referred to the template structure of AdpA (SMTL ID: 3w6v.1), and the amino acids are conserved in *Streptomyces* (Fig. 8b).

**Figure 8.**
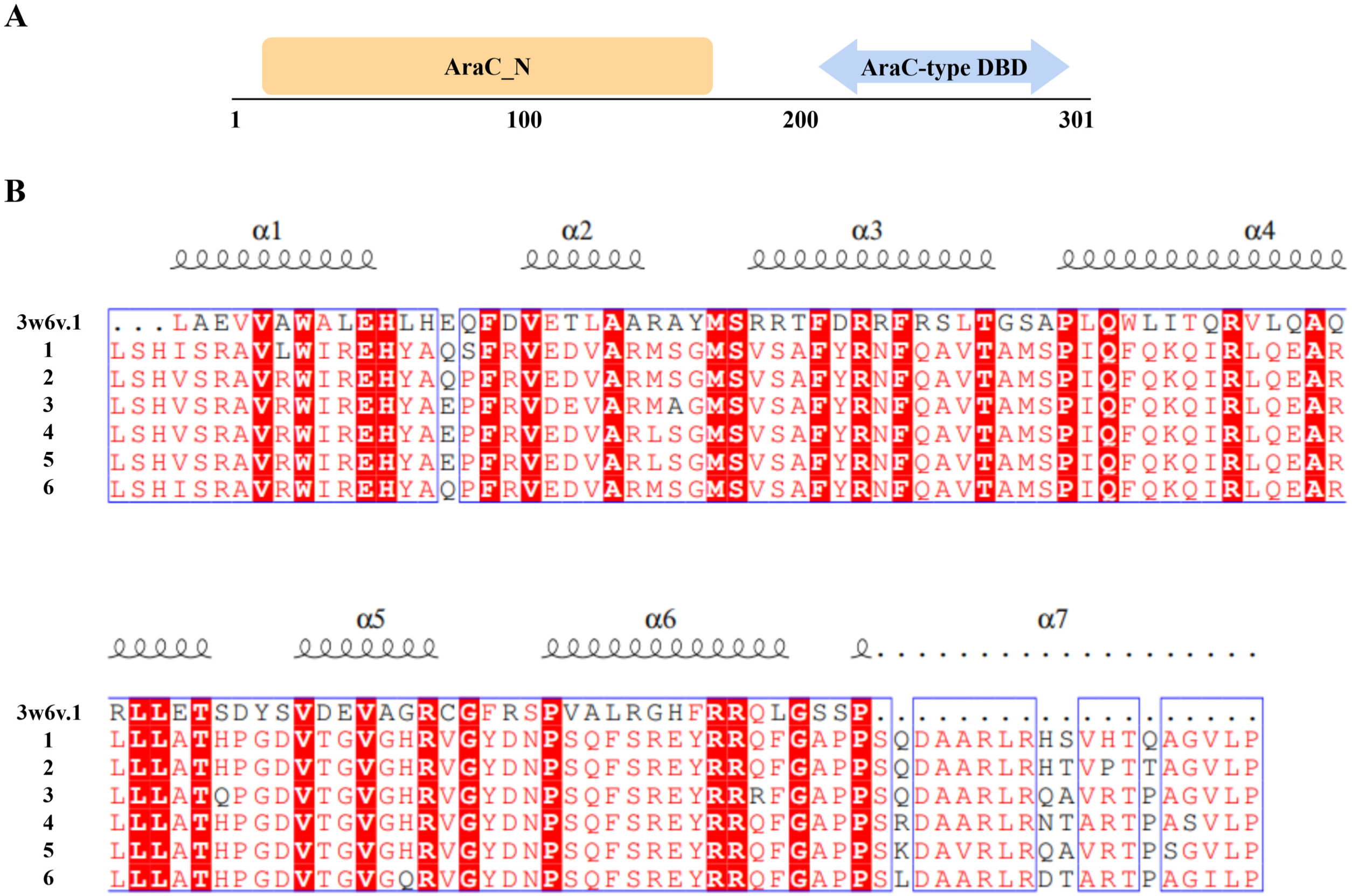
Functional domains and sequence alignment of SLINC_RS42780. **A** Predicted functional domains of SLINC_RS42780. **B** Alignment of the DBD domains of SLINC_RS42780 with related proteins. 3w6v.1, the template used for structure modeling of SLINC_RS42780 in SWISS-MODEL. 1, SLINC_RS42780 from *S. lincolnensis* (WP_067443797.1); 2, *Streptomyces albicerus* (WP_151477398.1); 3, *Streptomyces albiflavescens* (WP_189192684.1); 4, *Streptomyces canu*s (WP_059211104.1); 5, *Streptomyces davaonensis* (WP_015663102.1); 6, *Streptomyces scabichelini* (WP_165261112.1).

Then, a *SLINC_RS42780* disruption strain Δ*SLINC_RS42780* was constructed using the method as above. PCR products amplified by the primer pair JD42780-F/R were 2.8 kb and 2.2 kb with WT and Δ*SLINC_RS42780* as templates, respectively. A 2.5 kb band appeared only using pKCcas9d42780 as template, rather than WT or Δ*SLINC_RS42780* (Fig. 9a). These data indicated that Δ*SLINC_RS42780* was constructed successfully.

**Figure 9.**
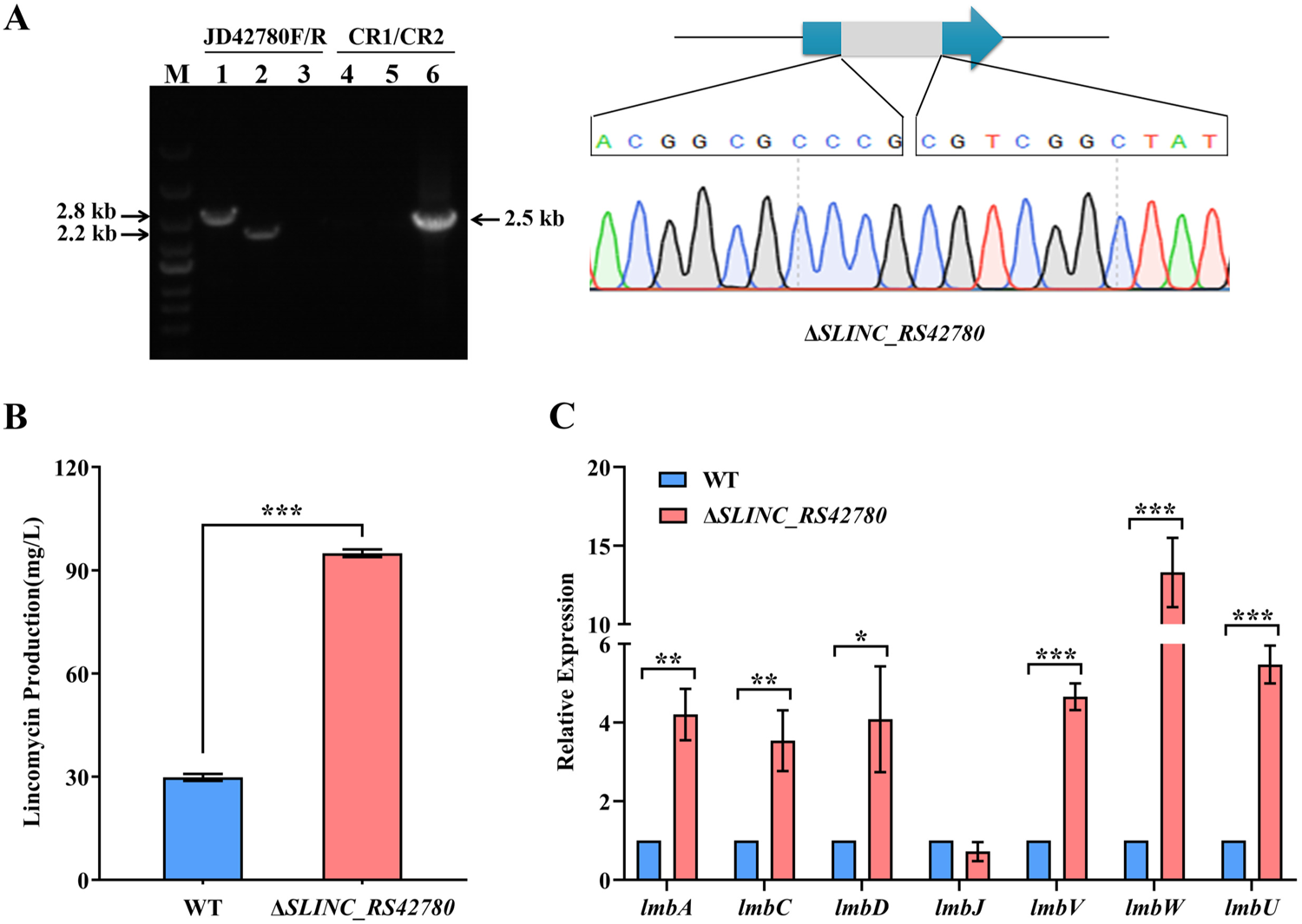
SLINC_RS42780 suppresses lincomycin biosynthesis. **A** Identification of Δ*SLINC_RS42780* by PCR and sequencing. Lane M indicated the DNA molecular weight marker. Lanes 1, 2 and 3 indicated PCR products amplified by primer pair JD42780F/R. Lanes 4, 5 and 6 indicated PCR products amplified by primer pair CR1/CR2. 1 and 4, WT; 2 and 5, Δ*SLINC_RS42780*; 3 and 6, pKCcas9d42780. **B** Effect of SLINC_RS42780 on lincomycin production. **C** Transcriptional analysis of lincomycin biosynthetic genes in WT and Δ*SLINC_RS42780*. The relative expression was normalized using internal reference gene *hrdB*. The transcriptional level of each gene in WT was set to 1.0. *, *P* < 0.05; **, *P* < 0.01; ***, *P* < 0.001.

Subsequently, the WT and Δ*SLINC_RS42780* strains were cultured in FM2 medium to measure the lincomycin production. The results of lincomycin bioassays showed that the yield of lincomycin in Δ*SLINC_RS42780* increased 3.2-fold compared to that in WT (Fig. 9b), indicating that SLINC_RS42780 negatively regulates lincomycin biosynthesis. Furthermore, qRT-PCR analysis demonstrated that, compared to WT, the transcriptional levels of *lmbA, lmbC, lmbD, lmbV, lmbW* and *lmbU* were significantly increased in Δ*SLINC_RS42780* with fold changes 4.2, 3.5, 4.1, 4.7, 13.3 and 5.5, respectively (Fig. 9c). The transcriptional levels of *lmbJ* were similar in WT and Δ*SLINC_RS42780*, suggesting *lmbJ* was not regulated by SLINC_RS42780. These data demonstrated that SLINC_RS42780 can suppress the transcription of *lmbA, lmbC, lmbD, lmbJ, lmbV, lmbW* and *lmbU*, thereby inhibit lincomycin biosynthesis.

### LmbU and its targets SLINC_RS02575, SLINC_RS05540 and SLINC_RS42780 coexist in many actinomycetes

Previously, we showed that LmbU homologues are widely found in actinomycetes, especially *Streptomyces* (39). It is very interesting to explore whether the regulatory targets of LmbU (SLINC_RS02575, SLINC_RS05540 and SLINC_RS42780) are also widely existed in actinomycetes. According to this consideration, we firstly searched the strains containing both the homologues of LmbU and one target in NCBI database. The results showed that all the homologues of SLINC_RS02575, SLINC_RS05540 and SLINC_RS42780 appear widely in various *Streptomyces* species which also contain LmbU homologues. The top 10 strains ranking by the identities of SLINC_RS02575, SLINC_RS05540 or SLINC_RS42780 were shown in Table S2, S3 or S4. Interestingly, some strains contain more than one LmbU homologues, such as *Streptomyces resistomycificus* (Table S2), *Streptomyces mirabilis* (Table S3), *Streptomyces albicerus* (Table S4), and so on. What’s more, a SLINC_RS42780 homologues was found in *Nonomuraea zeae*, which belongs to actinomycetes, but not *Streptomyces*, indicating the universality of LmbU and its regulatory targets.

In addition, the strains containing all the homologues of LmbU and the three targets (defined by identities more than 30%) were searched in NCBI database. The top 10 strains ranking by the identities of LmbU were shown in Table 3. The data revealed that all the four proteins are widely found in actinomycetes, including a non-*Streptomyces Kutzneria buriramensis*, indicating that LmbU homologues may also regulate these targets in other actinomycetes. Subsequently, we analyzed the promoter regions of the genes encoding the homologues of the three targets in the 10 strains. Surprisingly, only the promoter regions of the genes encoding SLINC_RS05540 homologue from *Streptomyces xylophagus*, and SLINC_RS42780 homologue from *Streptomyces albicerus* contain the 10-bp conserved palindromic sequence 5’-TCGCCGGCGA-3’ (Table S5). These data demonstrated that the regulatory mechanisms of LmbU homologues in different strains have similarities as well as differences.

**Table 3.**
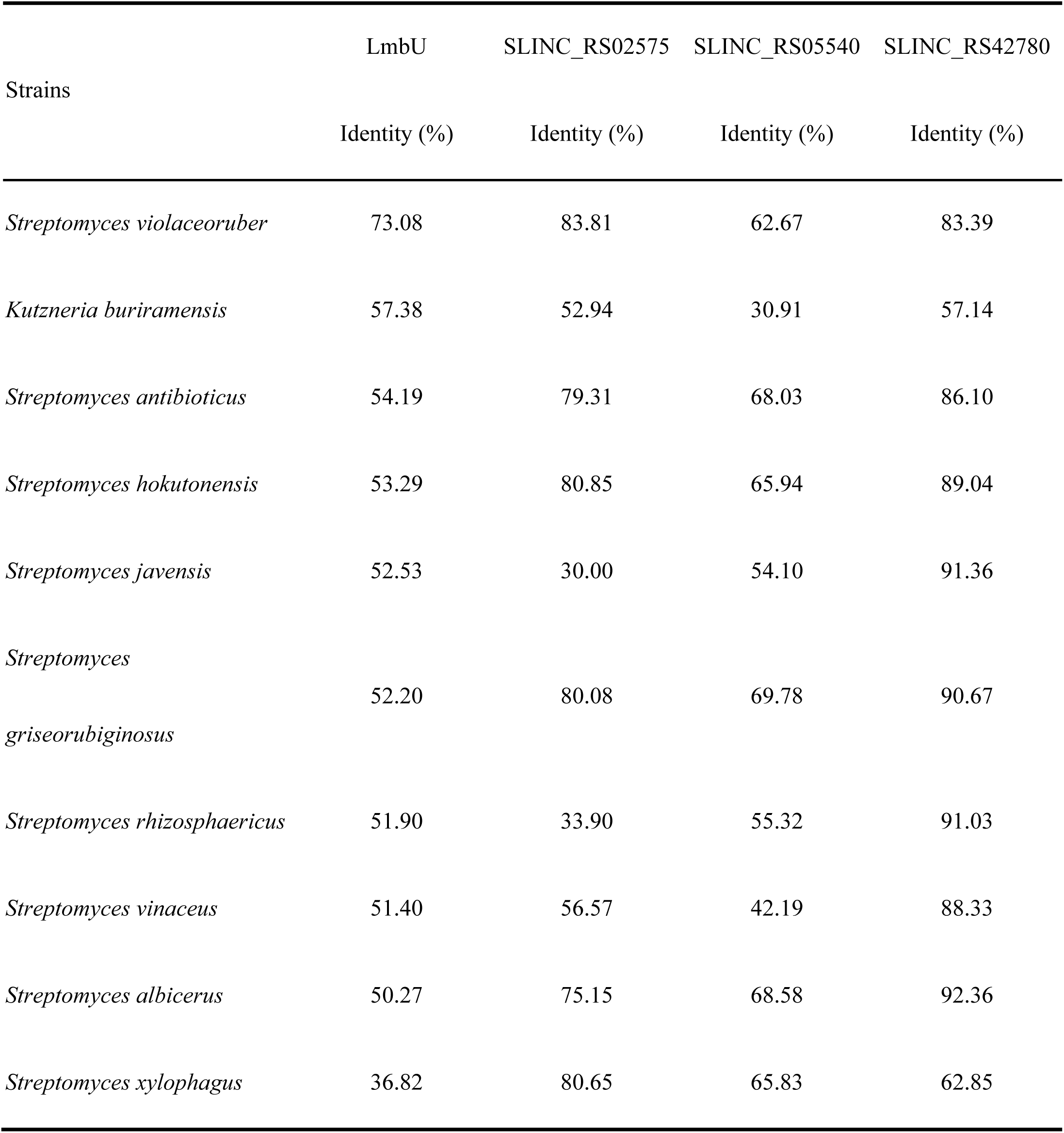
Strains contain all the homologues of LmbU, SLINC_RS02575, SLINC_RS05540 and SLINC_RS42780.

|  | LmbU | SLINC_RS02575 | SLINC_RS05540 | SLINC_RS42780 |
| --- | --- | --- | --- | --- |
| Strains | Identity (%) | Identity (%) | Identity (%) | Identity (%) |
| *Streptomyces violaceoruber* | 73.08 | 83.81 | 62.67 | 83.39 |
| *Kutzneria buriramensis* | 57.38 | 52.94 | 30.91 | 57.14 |
| *Streptomyces antibioticus* | 54.19 | 79.31 | 68.03 | 86.10 |
| *Streptomyces hokutonensis* | 53.29 | 80.85 | 65.94 | 89.04 |
| *Streptomyces javensis* | 52.53 | 30.00 | 54.10 | 91.36 |
| *Streptomyces*<br>*griseorubiginosus* | 52.20 | 80.08 | 69.78 | 90.67 |
| *Streptomyces rhizosphaericus* | 51.90 | 33.90 | 55.32 | 91.03 |
| *Streptomyces vinaceus* | 51.40 | 56.57 | 42.19 | 88.33 |
| *Streptomyces albicerus* | 50.27 | 75.15 | 68.58 | 92.36 |
| *Streptomyces xylophagus* | 36.82 | 80.65 | 65.83 | 62.85 |

## Discussion

LmbU, a CSR of lincomycin BGC, has been shown to directly or indirectly regulate the structural genes within the *lmb* cluster (25). In addition, we have demonstrated that LmbU homologues are widely found in actinomycetes, and their positions on the chromosome are not limited to the antibiotic BGCs (25, 39). Based on this, we screened and identified the targets of LmbU which are located outside the *lmb* cluster, and showed the effect of these targets on production of lincomycin (Fig. 10). LmbU promotes lincomycin biosynthesis through regulating transcription of the *lmb* genes as well as three target genes outside the *lmb* cluster. In addition, the three target genes, Δ*SLINC_RS02575*, Δ*SLINC_RS05540* and Δ*SLINC_RS42780*, have been found negatively regulated lincomycin biosynthesis via regulating transcription of the *lmb* genes including lmbU. However, it is unknown whether the three regulators directly regulate the *lmb* genes by binding to their promoters. This study can further illuminate the regulatory network of lincomycin biosynthesis, and will bring light to the functional analysis of LmbU family regulators. Cross-regulation of CSRs among disparate antibiotic biosynthetic pathways has been widely studied in *Streptomyces* (14-16, 20). However, it is rarely reported that CSRs regulate the targets outside the BGCs. Here, we found that the DBS of LmbU are widely distributed in the genome of *S. lincolnensis*, and 54 DBSs were located in the potential regulatory regions of genes, suggesting that LmbU is likely to function as a pleiotropic regulator and regulate more targets except for the *lmb* genes. We chose 14 targets which may be relevant to lincomycin biosynthesis, and carried out EMSAs. The results showed that LmbU can bind to 8 out of the 14 targets (Fig. 1), which encode regulators, transporters, sigma factors and other enzymes, indicating LmbU may function as a pleiotropic regulator. Though the sequence of DBS of LmbU within each target is perfectly matched, the binding affinity is not similar, indicating that the flanking sequence of DBS or the structure of the DNA may be also important for DNA-binding of LmbU. As reported, two ways have been found to be involved in DNA sequences recognition of proteins (42). One way is directly based on contacts of amino acids and bases, and the flanking sequences are also important (43). For example, the first and second flanking positions 5’ to the consensus DBS play important roles in DNA-binding affinity for E12 homodimer and E12-TAL1 heterodimer (44). The other way is indirectly mediated by the conformation of the DNA (45-47). Thus, the structures of DNA, including bendability, stability, groove shape, flexibility and so on, rather than the simple sequence are more appropriate to determine the DNA-binding affinities of proteins.

**Figure 10.**
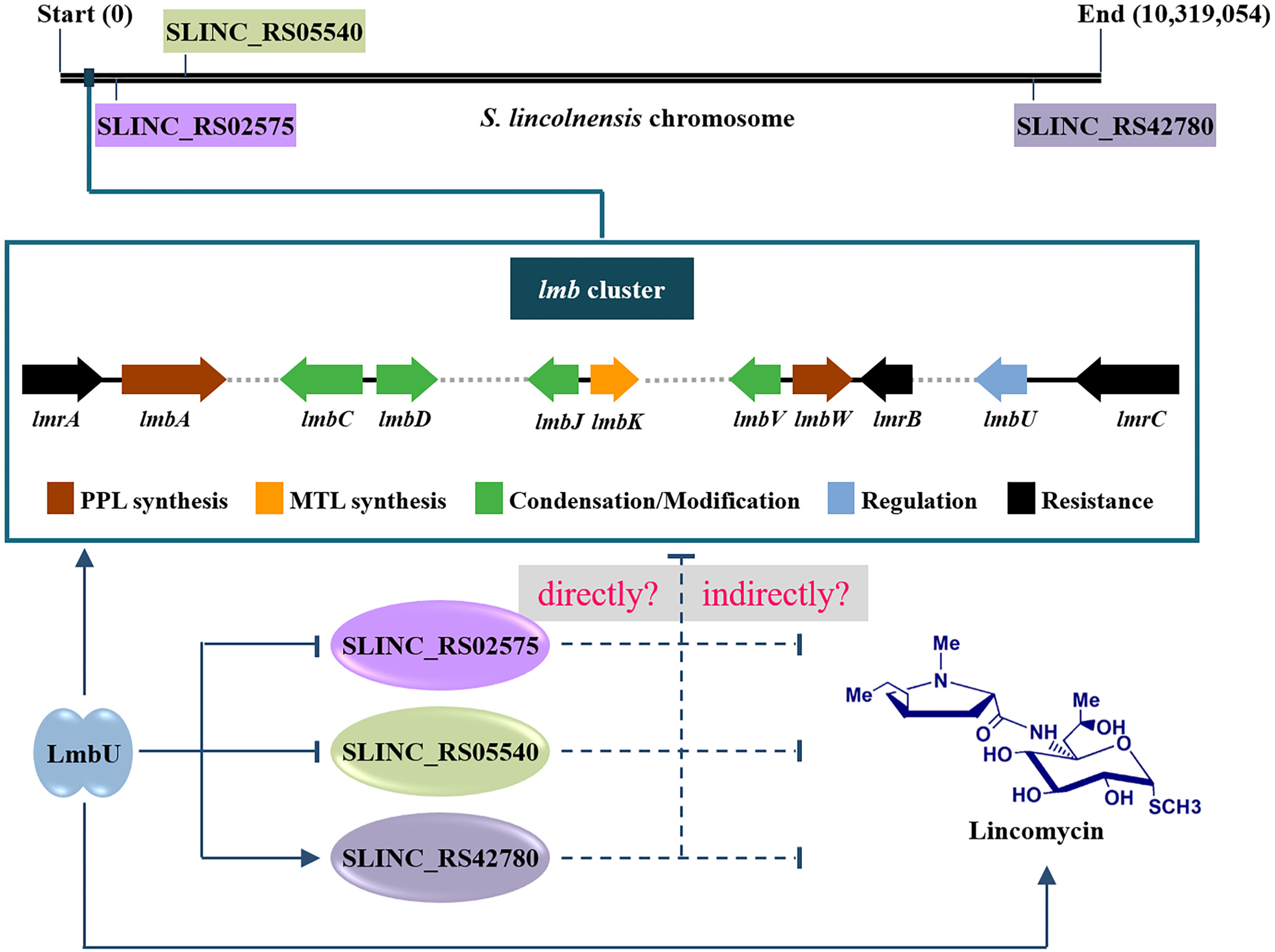
Proposed model of LmbU mediated regulation network to lincomycin biosynthesis. The locations of the *lmb* cluster and the three target genes on the chromosome were indicated. The arrows indicated activation, and the vertical virgules indicated inhibition. The solid lines indicated direct action, the dotted lines indicated unknown mechanisms.

Previously, we have showed that LmbU promotes the production of lincomycin by activating the transcription of *lmb* genes (25). Here, we demonstrated that LmbU inhibited the transcription of *SLINC_RS02575* and *SLINC_RS05540*, which negatively regulate the production of lincomycin, suggesting that LmbU affect the production of lincomycin not only by activating the *lmb* genes, but also by suppressing the genes against lincomycin biosynthesis. On the contrary, LmbU activates the transcription of *SLINC_RS42780*, which negatively regulates the production of lincomycin. This may be conducive to maintain the level of lincomycin within a certain range *in vivo*.

SLINC_RS02575 belongs to LAL family transcriptional regulator, which has both ATPase activity and DNA-binding activity. LAL family regulators generally function as CSRs and directly regulate antibiotics biosynthesis, such as SlnR for salinomycin biosynthesis in *S. albus* (48), RapH for rapamycin biosynthesis in *Streptomyces hygroscopicus* (49), and MilR for milbemycin biosynthesis in *Streptomyces bingchengensis* (50). SLINC_RS05540 belongs to AcoR family transcriptional regulator which contains an N-terminal GAF domain for signal sensing and ligand binding, a central AAA+ domain with ATPase activity, and a C-terminal DBD domain with a HTH motif associated with DNA-binding (51). AcoR has been well studied in *Bacillus subtilis*, and it functions as a σ^54^-dependent transcriptional activator (52) SLINC_RS05540 contains the C-terminal HTH motif and the central defective AAA+ domain, which includes the walker A motif, but lack of the walker B motif. In addition, the N-terminal of SLINC_RS05540 does not contain the GAF domain. Thus, we speculated that the functions of SLINC_RS05540 and AcoR may have commonality as well as diversity. SLINC_RS42780 belongs to AraC family transcriptional regulator, the members of which have been found in a variety of bacterial species and regulate the transcription of genes participated in carbon sources, stress responses, and so on (53). AdpA, a best studied member of AraC family, widely exists in *Streptomyces* and takes part in morphological differentiation and secondary metabolism (33). These data revealed that the regulatory network of LmbU on lincomycin biosynthesis is complex and accurate. In addition, LmbU can bind to the promoter regions of *SLINC_RS03185* (encode a MFS transporter), *SLINC_RS33920* (encode a sugar ABC transporter permease), *SLINC_RS38630* (encode a DHA2 family efflux MFS transporter permease subunit), *SLINC_RS34875* (encode a sigma 70 family RNA polymerase), and *SLINC_RS05745* (encode a methyltransferase domain-containing protein). The studies about the regulatory mechanisms of LmbU to these targets and the effects of these target genes on lincomycin biosynthesis are ongoing.

It is worth noting that LmbU homologues and SLINC_RS02575, SLINC_RS05540, SLINC_RS42780 homologues are all widely found in actinomycetes. After analysis of the 10 strains containing all the four proteins, we found that the promoter regions of only two genes contain the 10-bp conserved palindromic sequence 5’-TCGCCGGCGA-3’, while the locations of which in the two strains are different from that in *S. lincolnensis* (Table S5). These data demonstrated that the regulatory mechanisms of LmbU homologues are diverse in different strains, and our findings provide a basic study on the research of LmbU homologues.

## Acknowledgements

This work was supported by the National Natural Science Foundation of China (NSFS) (31900059), and the China Postdoctoral Science Foundation (2019M650079).

## Compliance with ethical standards

### Ethical approval

This article does not contain any studies with human participants or animals performed by any of the authors.

